# Accelerating the Characterization of Dynamic DNA Origami Devices with Deep Neural Networks

**DOI:** 10.1101/2023.05.11.540408

**Authors:** Yuchen Wang, Xin Jin, Carlos Castro

## Abstract

Mechanical characterization of dynamic DNA nanodevices is essential to facilitate their use in applications like molecular diagnostics, force sensing, and nanorobotics that rely on device reconfiguration and interactions with other materials. A common approach to evaluate the mechanical properties of dynamic DNA nanodevices is by quantifying conformational distributions, where the magnitude of fluctuations correlates to the stiffness. This is generally carried out through manual measurement from experimental images, which is a tedious process and a critical bottleneck in the characterization pipeline. While many tools to support analysis of static of molecular structures, there is a need for tools to facilitate the rapid characterization of dynamic DNA devices that undergo large conformational fluctuations. Here, we develop a data processing pipeline based on Deep Neural Networks (DNNs) to address this problem. The YOLOv5 and Resnet50 network architecture were used for the two key subtasks: particle detection and pose (i.e. conformation) estimation. We demonstrate effective network performance (F1 score of 0.85 in particle detection) and good agreement with experimental distributions with limited user input and small training sets. We also demonstrate this pipeline can be applied to multiple nanodevices, providing a robust approach for the rapid characterization of dynamic DNA devices.

## Introduction

Structural DNA nanotechnology is a rapidly growing field that has shown great utility in the bottom-up fabrication of devices and materials with applications spanning areas like nanofabrication [1], nanophotonics [2], molecular computation [3], bioimaging [4], and nanotherapeutics [5]. Over the last several years there has been a surge of interest in dynamic DNA nanotechnology, since the ability to design reconfigurable DNA nanodevices, combined with the ability to interface DNA with a wide range of biomolecules or nanomaterials, is highly attractive for the development of sensors [6], nanorobots [7], tunable plasmonic devices [8], and biophysical measurement tools [9]. Since many of these applications rely on physical interactions with other molecules or materials, understanding the mechanical properties of dynamic DNA structures is crucial to quantitively describe their functions. The most common approach to characterize the mechanical properties of dynamic DNA devices is through imaging (transmission electron microscopy (TEM) or atomic force microscopy (AFM)) to visualize conformational fluctuations. The magnitude of these fluctuations is related to the structure stiffness. However, quantifying these conformations is typically done through manual measurement, which is tedious and often the major bottleneck limiting the characterization pipeline and slowing down the experimentation and overall design and test cycle. Hence, there is a critical need for approaches that facilitate, and ideally automate, rapid characterization of structure conformations for a variety of dynamic DNA nanodevices.

Machine learning is a type of artificial intelligence that enables machines to learn to identify patterns from data and improve their performance on a specific task without being explicitly programmed. It involves the use of algorithms that can be trained (i.e., learn) to make predictions based on current observations. Among several different algorithms, the use of deep neural networks (DNN) has been the dominant approach for a wide range of data problems [10]. Specifically, efforts in both academia and industry [11] are using DNNs to solve challenging real-world problems such as autopilot [12]–[14], robotics [15]–[17], speech recognition [18]–[20], predictive analytics [21]–[24], and computer vision [25], [26]. For example, to date, AlphaFold, which is an algorithm based on DNN, has already provided over 200 million protein structures with high accuracy [27]. In contrast, from traditional X-ray crystallography or cryo-EM, there are only ∼200 thousand protein structures shared in the protein data bank (PDB). Specific to DNA nanotechnology, recent studies [28]–[30] showed the DNN is also a great solution for automatically recognizing nanostructure in atomic force microscopy or fluorescence microscopy with high accuracy, which provided a foundation to solving the DNA nanostructure identification problem. However, these works only demonstrated the feasibility of identifying static nanostructures from images and did not address the need for automated property characterization, which is necessary to overcome the characterization bottleneck for dynamic DNA nanodevices.

Here, we demonstrate a DNN pipeline that can accelerate the analysis of mechanical properties (i.e., flexibility) of dynamic DNA origami nanodevices [31], [32]. The pipeline implements two DNNs to facilitate the sampling (i.e., nanostructure identification) and quantification (i.e., conformation measurement) steps. We first establish the approach using a ‘Hinge’ nanostructure, which is representative of dynamic nanodevices that are widely used for biophysical measurements [9], biosensing [33], and controlling biomolecular interactions [34]. Secondly, we demonstrated the robustness and versatility of this pipeline by applying it to other dynamic DNA origami device characterizations including a ‘Hinge-Nucleosome’ system [9] and a three-arm devices designed to exhibit steric interactions between the arms [35]. Our results suggest that the DNN algorithm can be used to overcome the bottlenecks that require excessive labor work for post-processing of micrographs to characterize dynamic DNA nanodevices, which can greatly facilitate the design, experiment, and development cycle.

We will open the source of the dataset and the code of our pipeline after this paper being published.

## Results

### Dynamic DNA Origami Structure Analysis Workflow

We selected a DNA origami hinge structure [9] as the basis to develop our DNN pipeline, since hinges are simple dynamic devices that are widely used. In particular, we used a hinge that was recently demonstrated as a useful assay to measure the dynamic properties of biological samples [9] and apply high forces on nanometer scale [36]. This hinge structure (Figure 1) consists of two arms that are connected by several short single-stranded DNA (ssDNA) connections that form a hinge vertex. The two arms (∼70 nm in length) are highly stiff and can be regarded as solid bodies. The vertex is designed to be much more flexible, allowing for rotational motion of the two arms, which is primarily constrained to one degree of rotational freedom. The hinge exhibits preferred angular conformations; hence, it can be regarded effectively as a torsional spring. In order to apply these hinge devices (e.g., to apply forces to biomolecules [9], detect biomolecules [33], or control enzyme interactions [34]), it is critical to understand their mechanical properties. The mechanical property most relevant to the function of the hinge is the torsional stiffness. The torsional properties are typically considered in terms of a rotational free energy landscape, which can be determined from angular conformation distributions through Boltzmann inversion. Hence, a key step to characterizing the mechanical properties of hinge devices is measuring angular distributions.

**Figure 1.**
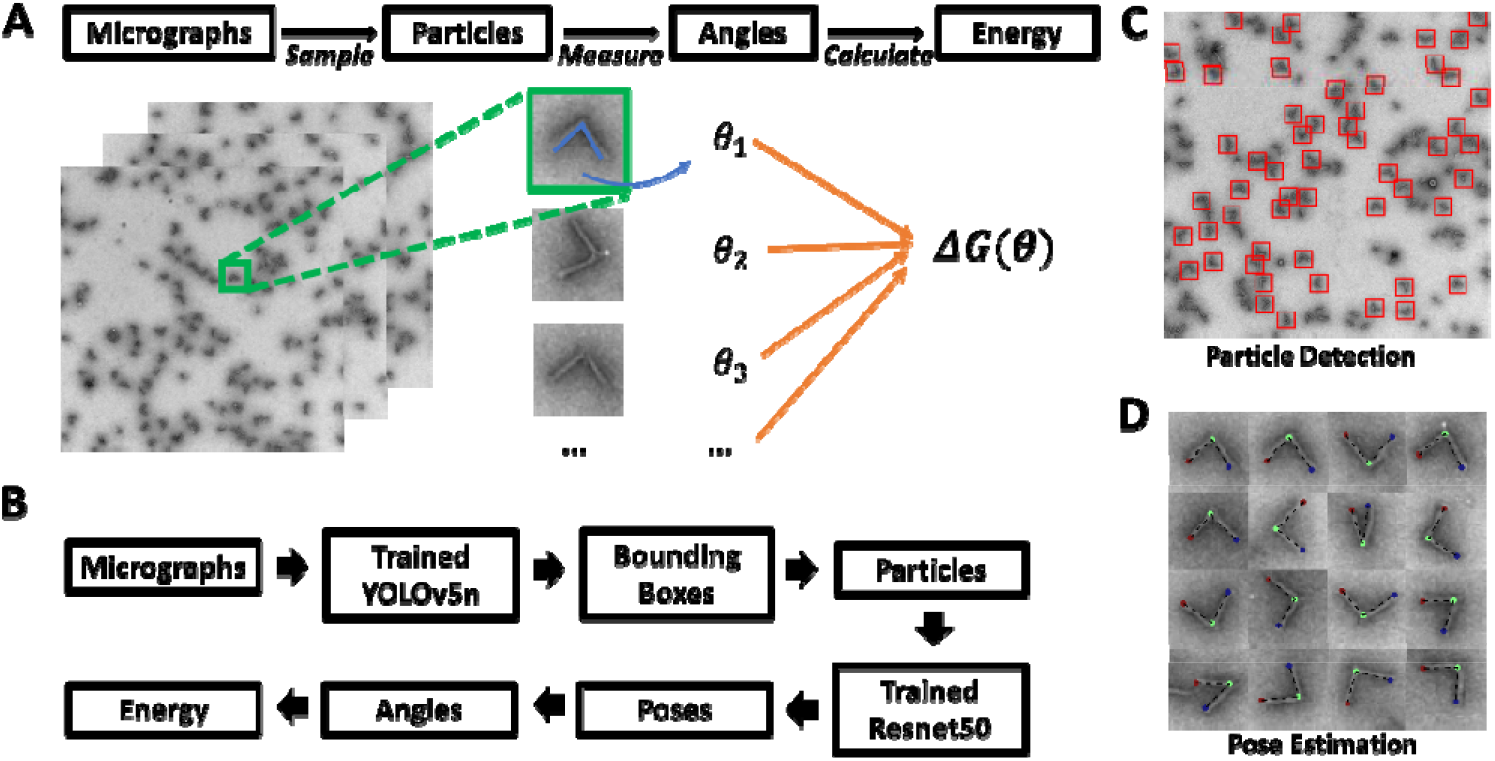
(A) The traditional pipeline for analyzing dynamic DNA origami device free energy landscapes with manual particle sampling and measurement. (B) The DNN-based pipeline. (C) Example micrograph illustrating particle detection. (D) Example particle montage illustrating pose estimation.

The most common approach to visualize the conformations of dynamic DNA origami devices is transmission electron microscopy (TEM) imaging [37], [38]. Most studies implement negative stain TEM, where dynamic nanodevices are deposited on a surface followed by imaging. The data analysis process to determine the conformational free energy landscape is typically carried out in multiple steps: 1) sampling many hinges (hundreds to thousands) by manual selection from images (i.e. clicking to identify a properly folded and isolated hinge nanostructure); 2) manually measuring their angles using tools like ImageJ [39]; and 3) determining an angular probability distribution from the manual measurement of hundreds to thousands of hinges; and 4) applying Boltzmann inversion to determine the angular free energy landscape [40]. This experiment and characterization pipeline is illustrated in Figure 1A. The manual sampling and measuring processes take a significant portion of time. For example, to determine one condition of free energy landscape, it includes ∼30 min of sample preparation, ∼30 min of imaging, but ∼2 hours for the manual characterization. Additionally, this amount of work can be easily scaled up by experiment iteration and repetition. Therefore, there is a clear need for a fully automated approach to accelerate the experiment cycle.

Hence, we introduce modified characterization pipeline that leverages DNNs to automate both the structure sampling and the conformation measurement. In our DNN-based pipeline, we utilize YOLOv5 network for sampling, where the DNN provides the center location and size (width and height) of a bounding box containing an isolated folded DNA origami hinge particle. Individual particle images were generated by cropping raw micrographs according to bounding boxes. Secondly, we utilize Resnet50 for angle measurements where it provides 3 critical positions with two hinge tips and one vertex points, as shown in Figure 1B. We refer to the first step as a ‘particle detection problem’ (Figure 1C) and the second step as a ‘pose estimation problem’ (Figure. 1D).

### Particle Detection Problem

We employed the DNN YOLOv5 [41] to solve our particle detection problem. There are several versions of the YOLOv5 network with various architectures and different complexities. Here, we used the smallest network YOLOv5nano (YOLOv5n. We also tested larger YOLOv5 networks, but they did not demonstrate significantly better performance, see Supplementary figure S2). In order to train the network, we first manually labeled (∼2-3 hours) the square bounding boxes from TEM micrographs for a total of 1257 individual particles from a total of 49 images as a ground truth reference. Based on the image index number, we then split the dataset (images + corresponding bounding box labels) into a training set (9 images), a validation set (10 images), and a test set (30 images). The motivation for the small training set is that we aimed to minimize the annotation work and computational cost to facilitate rapid training for future applications. The training set was used for fitting the YOLOv5n model. The validation set was used for tuning the network such as its architecture and hyperparameters to avoid overfitting. The test set was then used to evaluate the final performance of the trained network (See Methods for specific details).

As shown in Figure 2A, the raw TEM images have several different major features: 1) isolated hinges with two clear arms (target particles for sampling), 2) hinges with a different orientation (i.e., vertical orientation) where the hinge angle cannot be observed due to the TEM grid deposition, 3) local aggregation where two or more hinges are touching or very near each other, and 4) image background. The trained YOLOv5n was able to effectively identify the target hinges among all these features. In Figure 2A, each identified particle is labeled with its predicted bounding box. In the particle detection task with only single class (e.g., here is only hinge), the F1 score is typically used as the evaluation matrix. The F1 score is a balanced approach of precision value and recall value. Generally, the precision value is a measurement of false positive over all positive and the recall value is a measurement of how much of true sample are missed during prediction (see detailed definition in Method Section).

**Figure 2.**
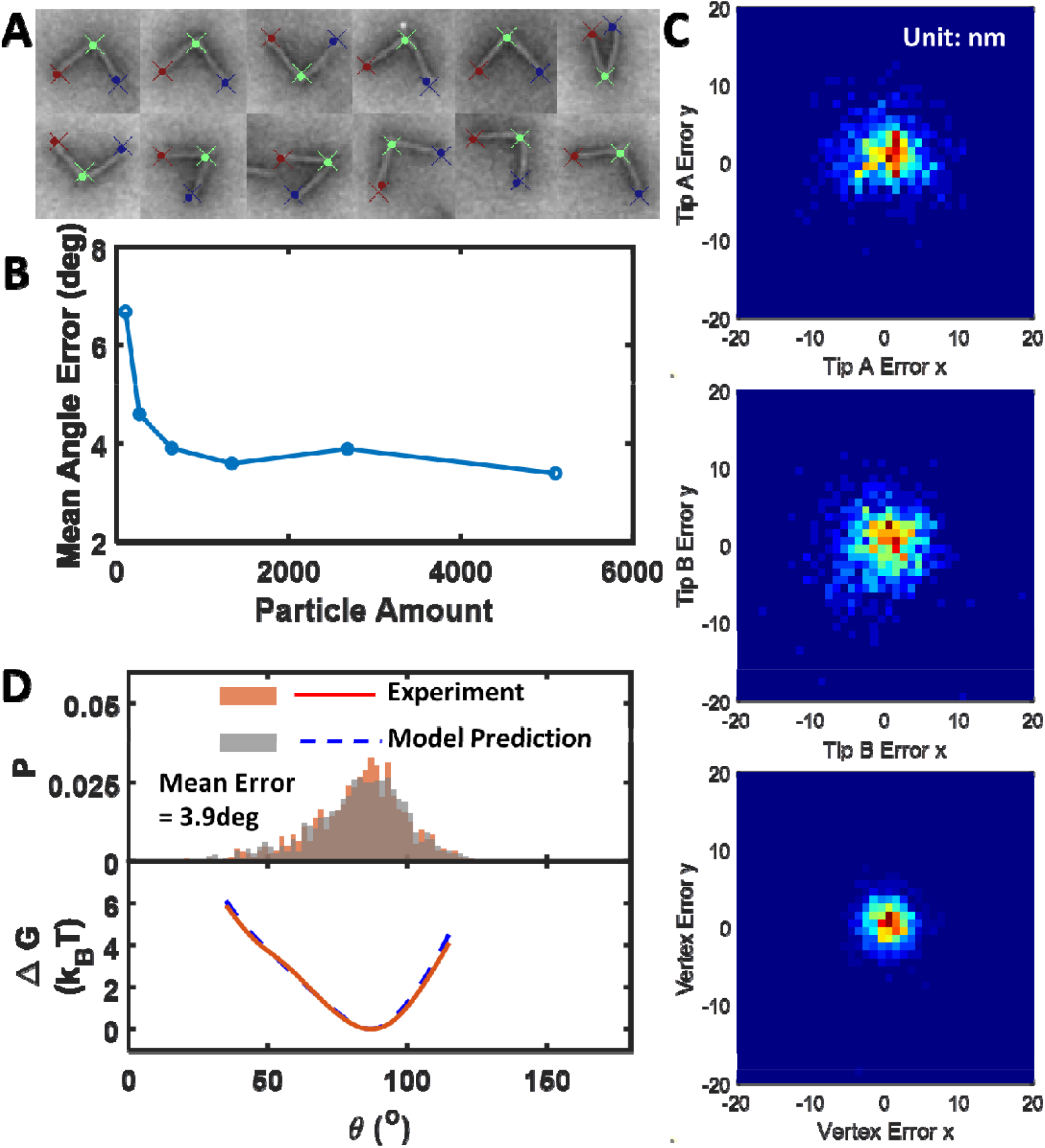
Pose estimation performance. (A) Random selected hinge particles with labels on. Cross: experiment. Dot: prediction. (B) Training Data size test (C) Spatial error distribution between prediction and experiment in the unit of nm (D) Downstream data test with hinge angle distribution and torque distribution.

Since we aimed to minimize the annotation labor work, we conducted training size sensitivity experiments by using a different number of TEM images for our training set. We found that only 6-10 images (∼30 target hinges per image) led to good network performance as indicated by the F1 score of ∼0.8 (Figure 2B).We selected 9 images since it gives highest F1 score and comparable labor work than 4-5 images.

To further improve performance, we also revisited the bounding box aspect ratio from prediction and found that many of the bounding box aspect ratios were not close to 1 as expected, especially for boxes on the image boundary. One potential way to mitigate this large aspect ratios would be to use a higher penalty value for the width and height loss, but this could influence the balance of bounding box position accuracy without careful tuning. By comparing with our ground truth annotation, we found these predicted boxes on the boundary with large aspect ratio contributed to a significant portion of false positives (Figure 2C). Therefore, we developed a bounding box-size filter (BBF) that 1) removes all boxes with greater than 1.5 aspect ratio; 2) re-defined the height and width of the bounding boxes to both be 50 pixels, which was the case for annotation. By doing so, we observed that the false positive value decreased from 217 to 141 (Figure 2D). This increased our precision from 0.83 to 0.88. We also noticed applying the BBF removed a minor fraction of true positives from 1033 to 1023, which reduced recall value from 0.82 to 0.81. Overall, the BFF increased the F1 score from 0.82 to 0.85 (summary of BFF effects shown in Figure 2D). Once the bounding boxes are defined by the network, all the predicted particles can rapidly be cropped out from the raw micrographs by image processing software such as MATLAB or Python.

**Figure 1.**
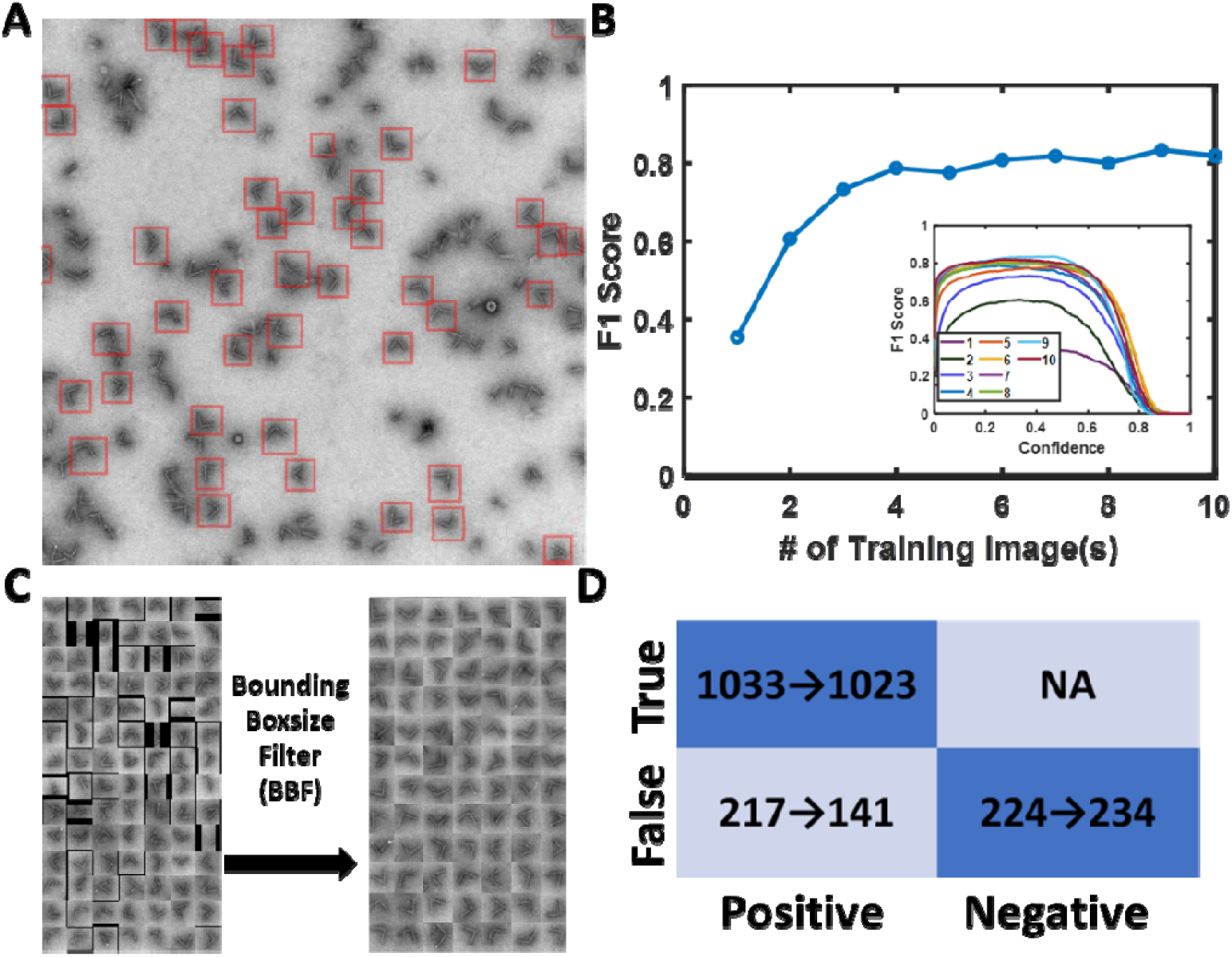
Particle detection performance. (A) An example of TEM image with YOLOv5 bounding box on the top. (B) Training data size sensitivity with F1 score. Inset, F1 score with different confidence value in variant training image number (C) The bounding boxsize filter removed particles with low and high dimensions. (D) Confusion matrix. the arrow represents the results after applying the filter.

### Pose Estimation Problem

To quantify the hinge angle conformation, we employed the Resnet50 neural network that was streamlined by DeepLabCut from Mathis Group [42]. To get a sufficiently large dataset for size sensitivity test, instead of only using the 1257 ground truth from labeled data in previous section, we also collected 5115 hinge particles from ref [9]. We manually annotated each particle with three critical points that define the hinge angle (two hinge tips A, B and one vertex that fit the inner lines of arms). We split the whole dataset into training (107, 269, 644, 1343, 2686, 5103 image particles), validation (269 image particles), and test set (1000 image particles).

To evaluate the sensitivity of training to the dataset size, we quantified the mean angle error for the Resnet50 model trained for different numbers of particles (Figure 3B). We found the error converged to below 4% when the training size was 644 particles or more, and even a training dataset size of ∼100 particles leads to lower than 8% angle error. For all evaluations in this section, we selected to use the network with training size 644. Figure 3C shows the spatial prediction error of the model compared with ground truth.

**Figure 3.**
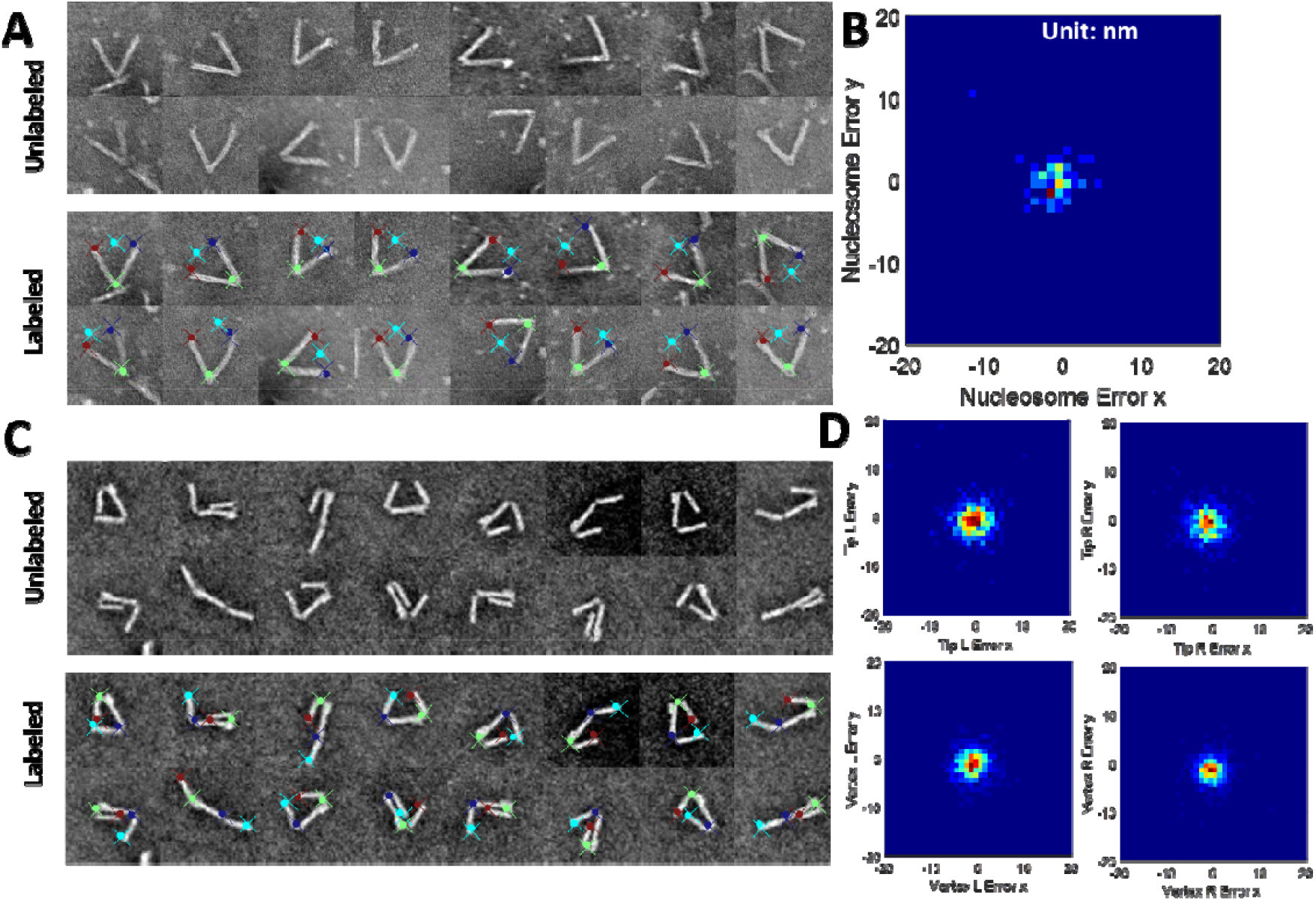
Pose estimation performance for Hinge-Nucleosome and SteriDyn dataset.

The hinge in TEM has no preference for orientation, and there is no specific feature difference between each tip. In other words, it is equivalent to arbitrarily flip Tip A and Tip B labels in the ground truth annotation. Therefore, it is seemingly impossible to consistently classify these datasets in the same order, and we likely have a ∼50% of flipped tip prediction due to randomness. If the misclassification happens, the spatial error would be significantly larger. but in fact we only have 3% flipped (Supplementary Figure S9). We assumed Resnet50 captured the spatial labeling pattern from the annotator, e.g., in Figure 3A, the blue tips are the first point, and the right tips are the third point during annotation. The blue points are higher than red points since the annotation was generally carried out from top to bottom on an individual particle.

We found the error distribution of tip A and tip B is larger than the vertex. We reasoned this was due to the different structural characteristics of the vertex and the tips. The vertex point was manually selected at the visually identified intersection of two lines along the inside of the arms; while tip A or tip B points were selected by visually identifying the tip along the inside of the arms, which is not as well-defined likely due to fraying of the ends. In practice, the tip position can be identified along the inner arm, but the distance away from the vertex is harder to define (see supplement Figure S10 for radial length error).

To further evaluate the capacity for the Resnet50 DNN to quantify mechanical properties, we converted the hinge angle measurements into angular probability distributions, and then calculated the free energy landscape from the probability distributions through Boltzmann inversion [40]. The free energy landscape gives a useful overall depiction of the relevant mechanical properties, and the applied torques and forces can be directly calculated from the free energy landscape as we have previously demonstrated [9]. We compared both the hinge angle probability distribution and angular free energy landscape predictions of the Resnet50 DNN to the experimental results (ground truth). Figure 3D shows a comparison of the angle distributions (top) and the angular free energy landscapes (bottom), illustrating the good agreement between the model prediction and the experimental results. Additionally, the two angle distributions passed the Two-sample Kolmogorov-Smirnov test (see Method for detail).

Ideally, the Resnet50 DNN pose estimation could be applied to a variety of dynamic DNA origami devices. To test the robustness of the approach, we applied the same Resnet50 architecture for two other previously obtained particle image datasets: 1) a dataset of hinge devices with incorporated nucleosome (i.e. DNA wrapped around a histone protein core, which is the base packing unit of genomic DNA in eukaryotes) where the nucleosome position is of interest [9], and 2) a dynamic devices with two fluctuating arms where the angular conformation of both arms are of interest [35]. Using the same training protocol, we trained Resnet50 models to predict specified critical features of the device conformations. In the first example, we are interested in quantifying the hinge angle, similar to the free hinge, and the nucleosome position as one additional coordinate point. Correlating the hinge angle and nucleosome position can be useful to study the wrapping/unwrapping of nucleosomes [9]. We labeled 321 image particles with 301 as training data. We found that the Resnet50 DNN can successfully recognize nucleosomes linked with hinge even in the presence of background noise such as free nucleosomes (white dots in Figure 4A). Figure 4B shows the error between the Resnet50 predicted nucleosome position and the manually determined position (ground truth), yielding an average error distance of 3.3 nm for nucleosome. In the second example we used a dynamic device with two fluctuating arms, which we refer to as the SteriDyn [35] due to the steric dynamic interactions of the arms. In this example, we are interested in the conformation of two moving components on a single device (i.e., the angle of each arm relative to the base platform). We manually annotated 4469 particles with 3500 as training dataset from TEM images using four points to define the two hinge angles. Figure 4C shows Resnet50 DNN can successfully recognize the orientation of the structure even though the left arm and right arm are very similar. The average error distances are 4.6, 4.5, 5.5, 4.8 nm for tip L, tip R, vertex L, vertex R, respectively.

## Discussion

Here, we demonstrated a DNN-based pipeline for quantifying the structure and mechanical properties (i.e., free energy landscape) of dynamic DNA origami devices using DNNs to automate two key steps of the characterization, namely particle detection and pose estimation (i.e., conformation measurement). We divided this workflow into two parts: 1) using a YOLOv5 DNN to detect the DNA origami structures from raw TEM images, and 2) then using a Resnet50 DNN to detect the conformation from individual particle images. In the particle detection process, we used a small training set (9 TEM images led to the highest F1 score in the current study) and simple network complexity. We further used BBF filters to remove a set of false positives that mostly happened at the image boundary and increased F1 score from 0.82 to 0.85. In the pose estimation process, we used 644 particles as a training set and achieved 3.9 deg hinge angle mean error. Furthermore, we demonstrated that our pose estimation process worked well for two other dynamic DNA nanodevices: Hinge-Nucleosome and SteriDyn, both of which agreed well with experimental data. The Hinge-Nucleosome example shows the pose estimation can be useful for devices applied to probe a molecule or interaction, and the SteriDyn example shows the pose estimation can be useful for systems containing more than one dynamic component/feature of interest.

The DNNs presented here were developed as a toolbox for finding patterns and quantifying specific features from experimental data. While the DNNs can perform forward evaluation within seconds, which is much faster than the manual approach, it is important to note there is significant manual effort required for the annotation of datasets (training, validation, and test datasets). Limiting this laborious process is the motivation for using small datasets to train the network. Therefore, the time saved for the entire experiment labor work would be dependent on the target throughput. For example, it typically takes more than 300 particles for sampling dynamic structures with a single condition [9], [35], [36]. Doing the manual annotation for a dataset of ∼300-500 particles to directly provide the ground truth could be completed within several hours, which is comparable to the time required to obtain datasets for training and testing DNN performance. Hence, developing an automated pipeline may not be worthwhile for a single dataset. However, in practice, dynamic DNA origami designs often take multiple iterations for optimization, or it may be of interest to design several versions with distinct properties (i.e., distinct conformational probability distributions and free energy landscapes). This often leads to anywhere from several to tens of datasets that need to be analyzed where the automated pipeline can make a major difference. As long as these cases involve the same basic structure with the same basic features, it should be possible to apply the same DNN pipeline, saving days or even weeks’ worth of manual annotation.

Rather than pursuing higher model precision, we sought to balance the amount of annotation work while providing effective results for the data analysis. For example, our results showed our pipeline led to 3.9 degree errors angle measurements and passed the Two-sample Kolmogorov-Smirnov test. Nevertheless, there are multiple ways to improve the network performance in terms of precision such as hyperparameter tunning, dataset redistribution, network architecture modification. For example, users can simply add more training images for better performance. However, the labor work could significantly increase, and this may conflict with the purpose of using DNN in this work.

More broadly, this general workflow can not only solve the mechanical properties of dynamic DNA nanodevices, but also could be suitable for non-DNA materials such as antibodies or other protein complex structures that undergo significant thermal fluctuations or conformational changes.

## Methods

### Preparation of the DNA Origami Structures

The DNA structures used in this work are based on scaffold DNA origami, which consists of a long single-stranded DNA (ssDNA) scaffold (M13MP18 bacteriophage virus prepared in our laboratory as described in [43]) and ∼200 short ssDNA staples. Based on Watson-Crick base-pairing rules, the design of staples sequences determines the assembly of hinge and SteriDyn structures, which are both previously reported [9], [35]. All staples were ordered from a commercial vendor (IDT, Coralville, IA). In the experiment, a final concentration of 20 nM scaffold, 200 nM of each staple strand,5 mM Tris, 5 mM NaCl, 1 mM EDTA, and 18 mM MgCl_2_, at pH 8.0 in aqueous solutions were made and then subjected to thermal annealing in a thermal cycler (Bio-Rad, Hercules, CA) for self-assembly. After that, the excess staples were purified by centrifugation in a polyethylene glycol (PEG) solution [44]. The remaining structures were resuspended with buffer (0.5xTBE with net 10mM MgCl_2_) and quantified by NanoDrop (NanoDrop 2000C Spectrophotometer, Thermo Scientific). The structures were diluted to 1 nM for downstream imaging.

### TEM Imaging

Transmission electron microscopy (TEM) was used to visualize the structure with nanometer resolution. Specifically, 4 μL of sample droplet was deposited on Formvar-coated copper TEM grids, stabilized with evaporated carbon film (Ted Pella; Redding, CA) for 4 min. The droplet was wicked away by filter paper and then stained by applying 7 μL 2% uranyl formate (SPI, West Chester, PA) and wicked away twice for 2 seconds and 15 seconds, respectively. TEM imaging was carried out at the OSU Campus Microscopy and Imaging Facility on an FEI Tecnai G2 Spirit TEM at an acceleration voltage of 80 kV at a magnification of 45,000x.

### Particle Detection Process with YOLOv5

The 50 raw TEM images that we used for training, validation, and testing were resized to 960×960 pixel jpeg files and further split into a training set (10 or less, we selected 9 images as the final training set), a validation set (10 images), and a test set (30 images). The manual annotation work was conducted in Roboflow [45] and modified by custom MATLAB scripts for adjusting all bounding boxes with a 50×50 pixel size. The prepared dataset was augmented stochastically by using the YOLOv5 default value(hyp.scratch-low.yaml). The neural network started from a pre-trained model (yolov5n) and it took ∼ 5 minutes on a machine with a RTX3060 graphics card RTX3060 (∼12 minutes on 1080Ti) for 500 epochs with the converged loss function. Specifically, YOLOv5 provided the x-position, y-position, width, height, and confidence of the predicted bounding box. The F1 score value was evaluated with the validation set for determining the optimal network parameters. After YOLOv5n network was selected, the test set was used for evaluating F1 score. The confidence threshold and IoU are selected to 0.47, and 0.3,respectively. The bound boxsize filter (BBF) was then employed for improve F1 score by removing bounding box with lower than 1.5 aspect ratio.

The precision (Pr), recall (Re), and F1 score are defined as:

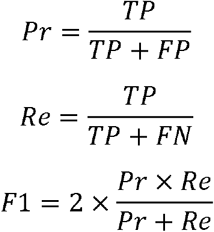

where TP, FP, FN represent true positive, false positive, false negative, respectively.

### Pose estimation process with Resnet50

All particles in this work were resized to 200×200 pixel jpeg files and split into a training set (644 particles), a validation set (4728 particles), and a test set (1000 particles). The manual annotation work was conducted in ImageJ [39]. Specifically, we used the ‘angle tool’ to mark two hinge tips and one vertex for each particle and the encrypted ROI data was parsed to xy position csv files using custom MATLAB scripts. The dataset was processed by the pose-estimation tool DeepLabCut [42]. By default, we used ‘imgaug’ image augmenter and the Resnet50 network. This neural network was finetuned from a pre-trained model ‘ImageNet’ and it took ∼25 min (on RTX3060) for 50k iterations to converge. The confidence threshold was selected to 0.92 to eliminate the majority of higher pixel errors. The performance was evaluated by pixel error, absolute angle error, and hinge energy. For Hinge-Nucleosome, the training size is 304 and the test size is 76. For SteriDyn, the training size is 3500 and the test size is 875. All others methods were identical compared with hinge structure.

### Statistical Test

To quantify the agreement of angle distribution between experiment and network prediction, we gave a Kolmogorov–Smirnov (KS) test. The null hypothesis is that two angle arrays are from the same continuous distribution. We tested these two angle arrays with build-in ‘kstest2’ function in MATLAB [46]. The p value [47] is 0.48 and is much greater than the significance level is 0.05. Therefore, this indicates a failure to reject the null hypothesis, which states network predictions are in a great agreement with experiment.

## Acknowledgements

This work was supported by the National Science Foundation through grants CMMI-1921881 and EFMA-1933344. We would like to thank all members of the Castro Lab and Soghrati Lab for helpful discussions and feedback. Transmission electron microscopy images were acquired at the OSU Campus Microscopy and Imaging Facility, which is supported in part by grant number P30 CA016058, National Cancer Institute, Bethesda, MD.

## Author contributions

Y.W and C.E.C. conceptualized the project, guided the execution of experiments, and supervised the research team. Y.W. led the execution of experiments including dataset preparation, network training, network evaluation. Y.W. led the MATLAB and Python code that were written for implementation. X. J. supported the experimental work and provided the critical network feedback. Y.W. and X. J. led the initial drafting with all authors providing editing and feedback. C.E.C. acquired the funding for this work. All authors have read and agreed to the published version of the manuscript.

## Additional information

Supplementary Information The online version contains supplementary material available at XXXXX.

