## Supplementary Information for "Accelerating the Characterization of Dynamic DNA Origami Devices with Deep Neural Networks"

**Supporting Information:**

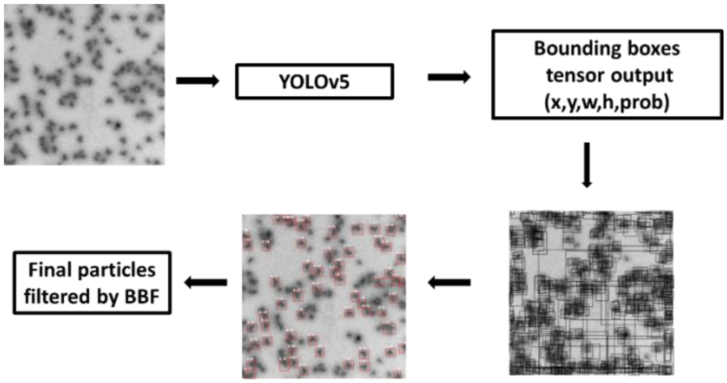


Figure S1 Generalized 'particle detection problem' input-output inference workflow. The images are processed by YOLOv5 after multi-stage convolutional operations. The output of network are a set of grids with each box has x, y, w, h, probability, respectively. The data on the grid was converted into bounding box. The probability value was used as threshold to balance the network precision and recall. See https://github.com/ultralytics/yolov5 for details. Finally, the BBF was applied to further improve the network performance.

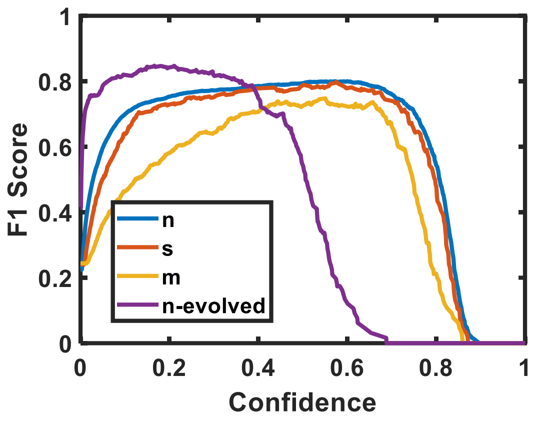


Figure S2 The performance comparison between several models provided by YOLOv5 with different complexities. n: nano, s: small, m: medium. Evolved represents a model trained by using an optimized hyperparameter from the genetic algorithm. Training size =10 images.

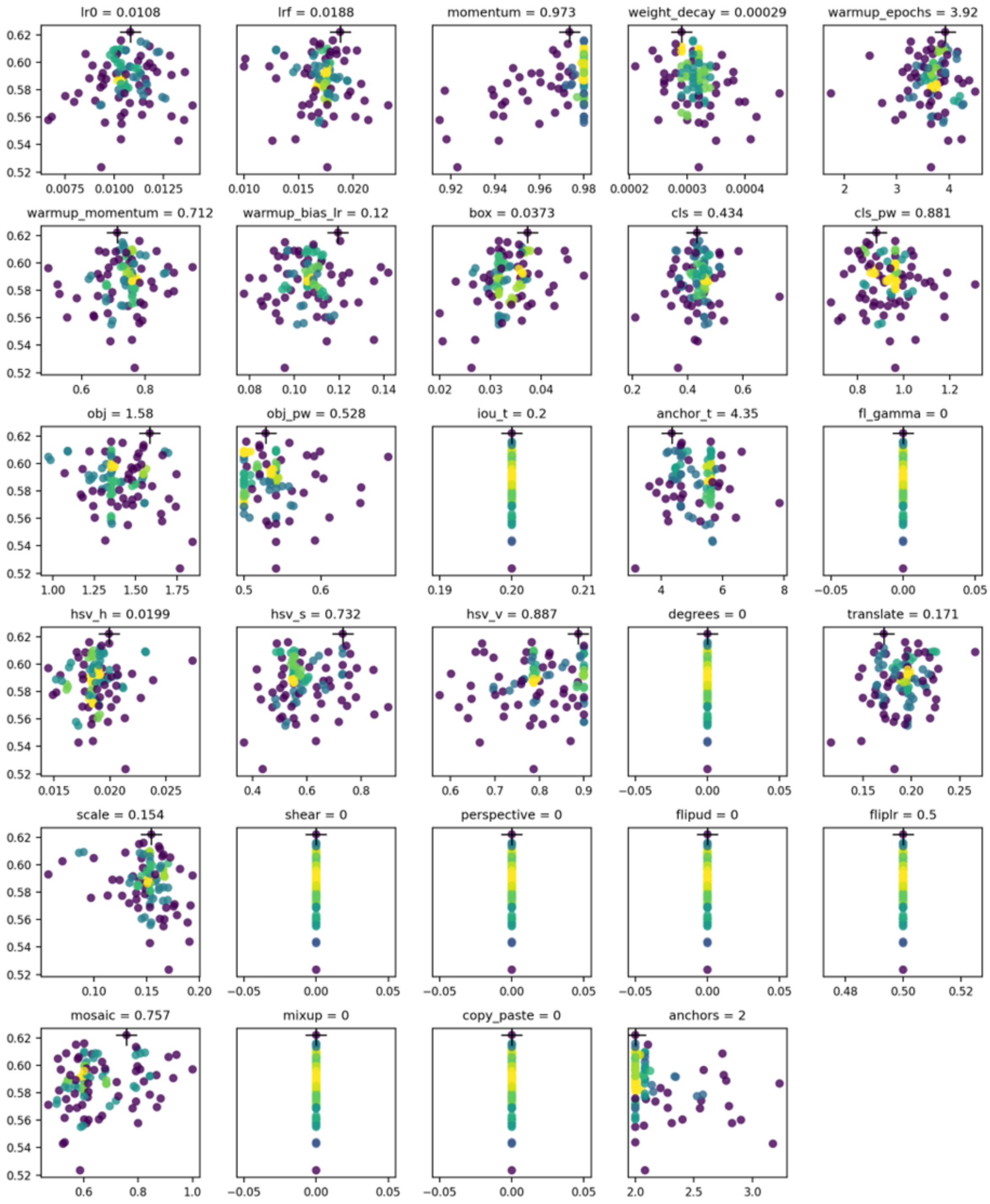


Figure S3 Optimization of hyperparameters matrix by genetic algorithm. The X axis is the hyperparameter value and the Y axis is the fitness value. The fitness function was defined as f=0.1*(mAP@0.5) +0.9*(mAP@0.5:0.95). Color represents dots density. The final hyperparameters were selected with the highest fitness after 300 generations with 500 epochs of each.

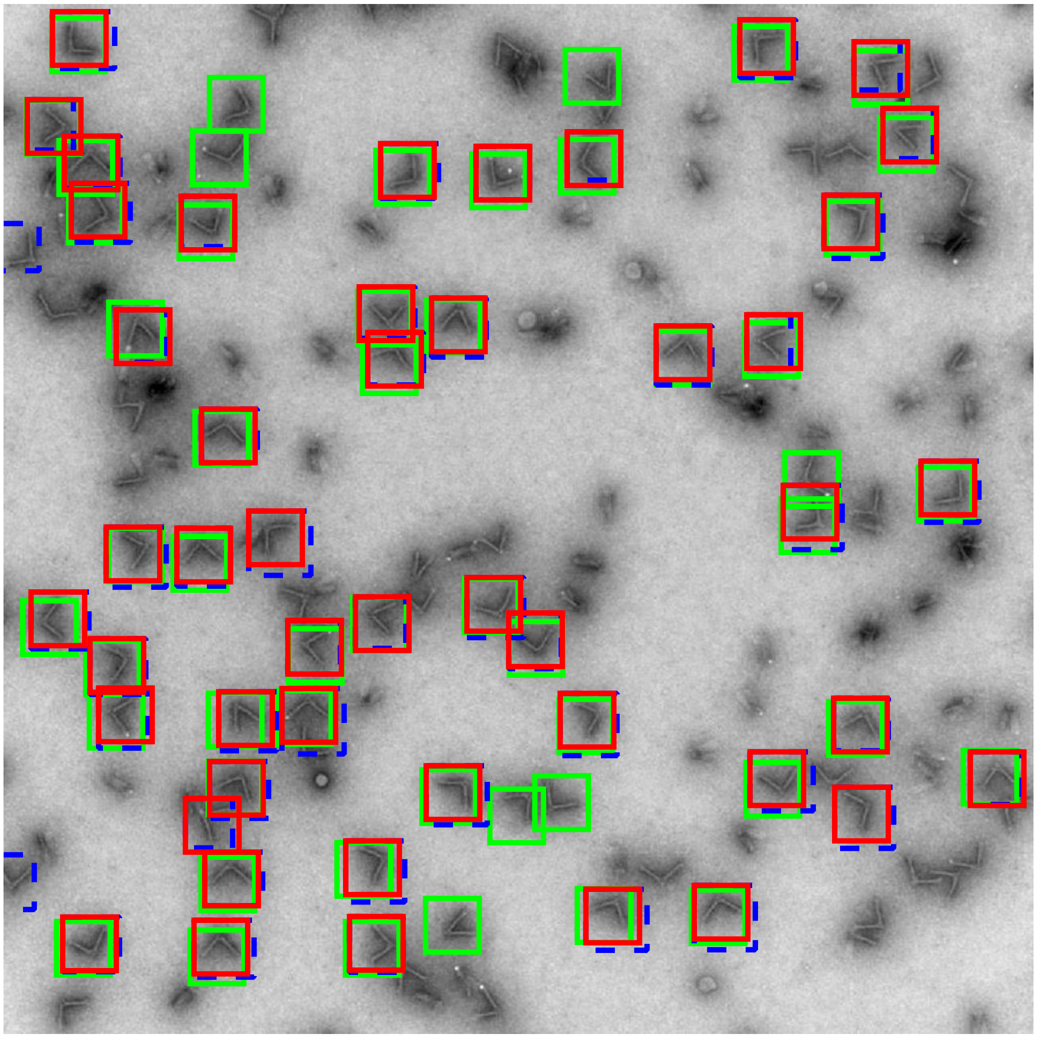

Figure S4 Comparison between ground truth (green), prediction for YOLOv5 (blue), and prediction after BBF (red). The blue-red offset effect is due to the resize of bounding box from predicted weigh and height into 50x50 pixel.

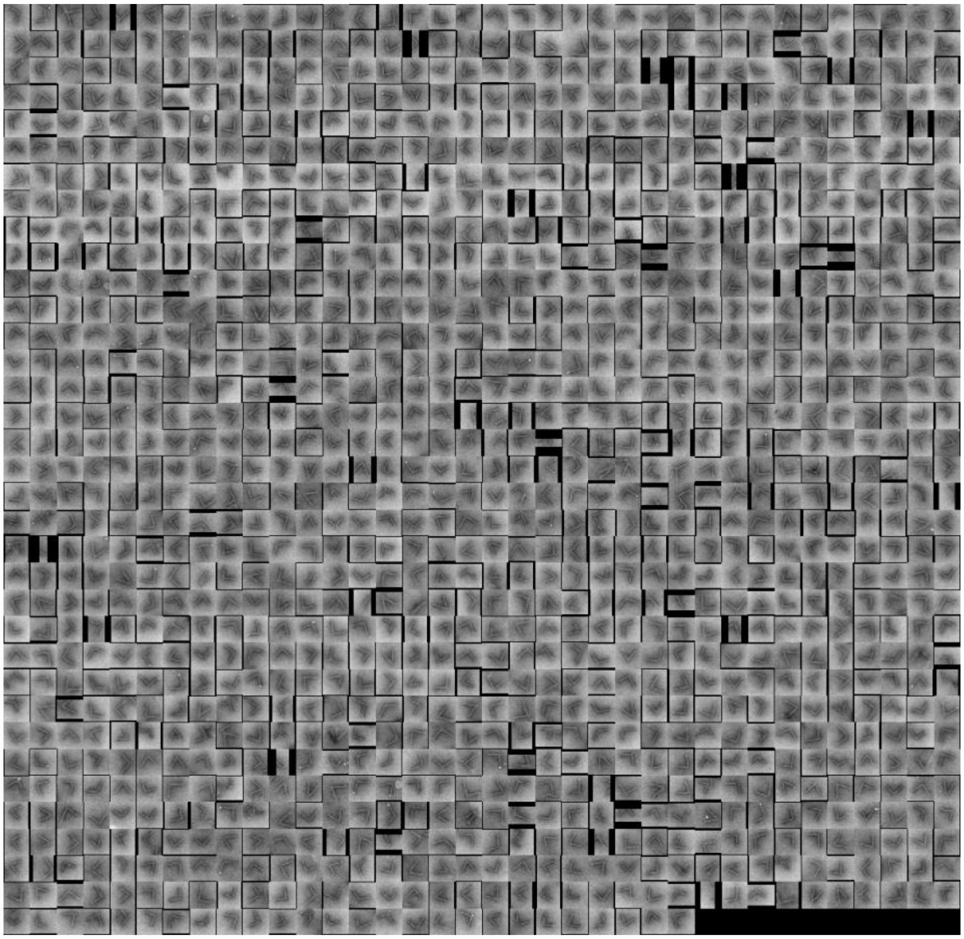

Figure S5 A gallery collection of predicted hinges from YOLOv5.

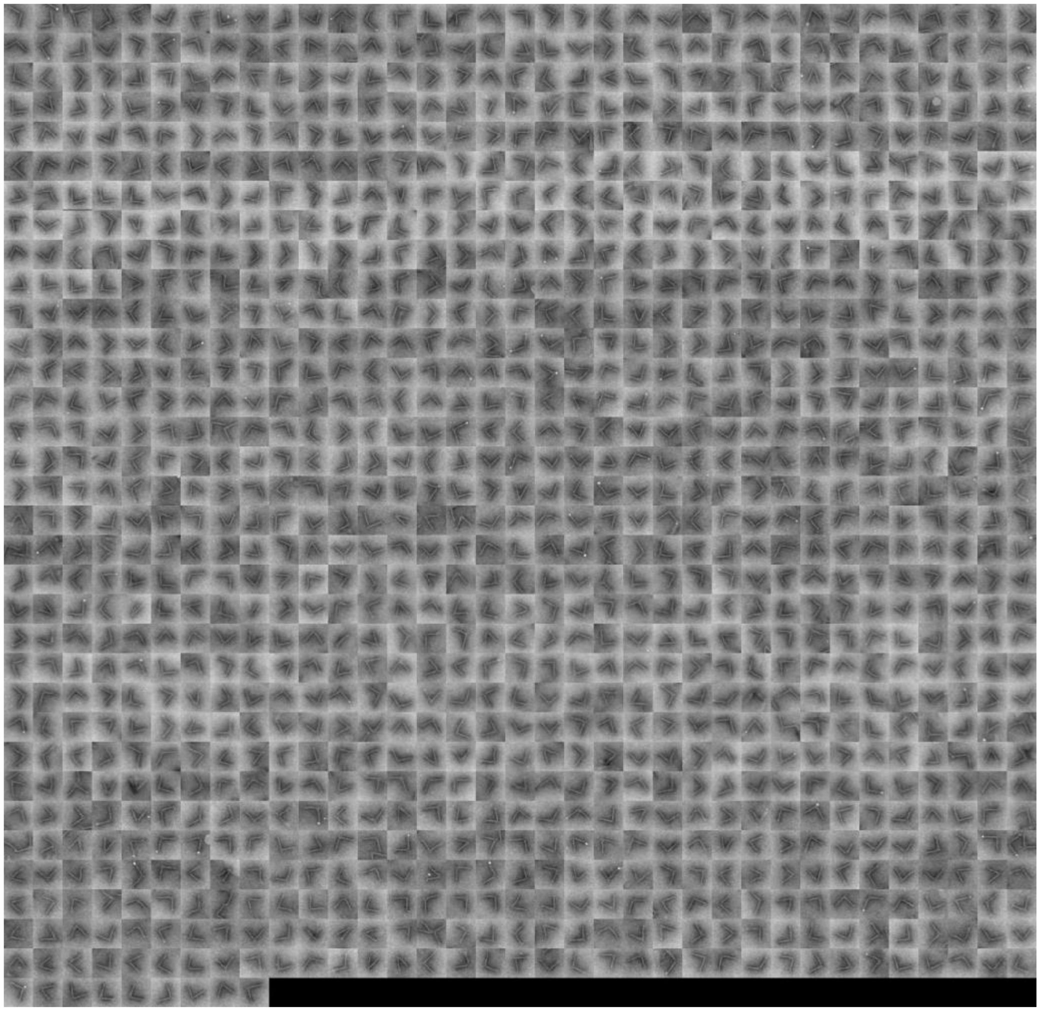

Figure S6 A gallery collection of predicted from YOLOv5 after BBF.

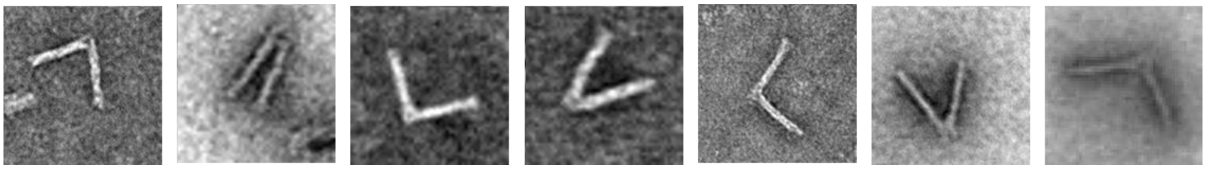

Figure S7 Heavy metal Uranyl formate staining process to TEM grid gives random image contrast behavior.

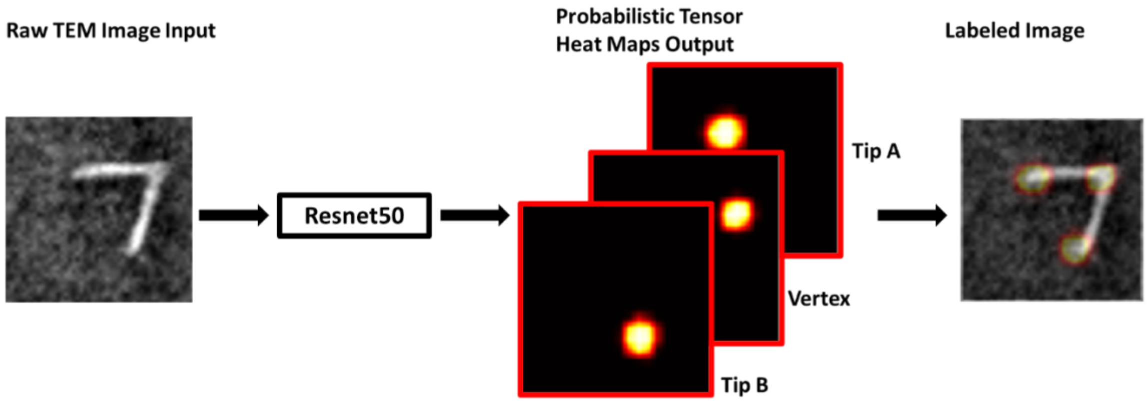


Figure S8 Forward evaluation process to determine the pose in Resnet50.

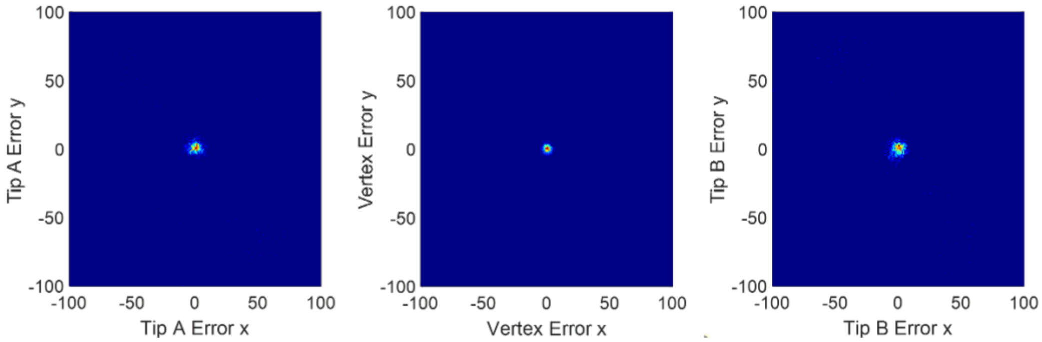


Figure S9 Hinge pose error in a large axis limit.

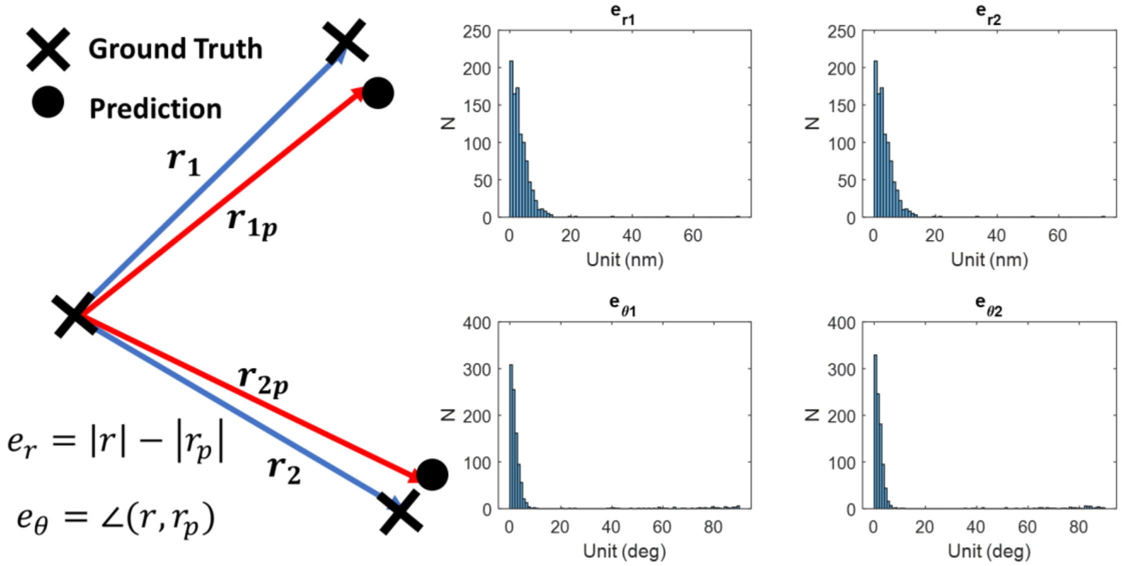


Figure S10 Hinge pose estimation error in term of radial length and theta.
